# Limited genetic variation for male mating success reveals low evolutionary potential for thermal plasticity in *Drosophila melanogaster*

**DOI:** 10.1101/166801

**Authors:** John T. Waller, Anna Kell, Mireia Ballesta, Aude Giraud, Jessica K. Abbott, Erik I. Svensson

**Author notes:** Data is archived on https://datadryad.org/.

## Abstract

Populations respond to novel environmental challenges either through genetic changes, through adaptive phenotypic plasticity for the traits in question, or by a combination of these factors. Here, we investigated the evolutionary potential of phenotypic plasticity for male mating success, locomotory ability, and heating rate (a physiological performance trait) in the fruitfly *Drosophila melanogaster*, using isogenic male lines from the Drosophila Reference Genome Panel (DGRP) and hemi-clonal males. We quantified thermal reaction norms of how male mating success changed in relation to a temperate gradient, ranging from cold (18 °C) via optimal (24 °C) to hot and stressful environments (either 30 °C or 36 °C). We found significant differences in male mating success and locomotory performance between different lines, as well as significant main effects of temperature, but no significant genotype-by-environment interactions (GEI:s). A statistical power analysis revealed that the variance explained by GEI:s for thermal plasticity using this sample size is likely to be modest or very small, and represent only 4% of the total variation in male mating success. The lack of strong GEI:s for these two behavioral traits contrast with the presence of significant GEI:s for male heating rate, as measured by thermal imaging (infrared camera technology). These results suggest that sexual selection through male mating success is not likely to be efficient in mediating evolutionary rescue through changed plasticity in response to changing temperatures.

## Introduction

Populations can respond to environmental challenges either evolutionarily through changes in allele frequencies or through adjustments in phenotypic plasticity (Schlichting and Pigliccui 1998). Phenotypic plasticity is the capacity of a single genotype to change its phenotype under different environmental conditions (Bradshaw 1965; Roff 1997; Pigliucci 2001). Phenotypic plasticity can increase a population’s mean fitness across several environments, and plasticity might increase niche space and geographical range (Ayrinhac et al. 2004; Manenti *et al*. 2015; Mather and Schmidt 2017). Responding plastically to changing environmental conditions is, however, a short-term survival strategy for a population, as there are costs and limits to plasticity (Lande 2014; Murren et al. 2015; Sgro et al. 2016), which can limit the potential to respond to sustained environmental change (Gonzalez et al. 2013).

Adaptive evolution of phenotypic plasticity requires not only that a population responds to changing environmental conditions, but the different genotypes must also differ in how they respond to these changing environmental conditions (Lande 2009; 2014; Chevin et al. 2010). A population must thus harbor enough standing genetic variation in environmental reaction norms to respond to sustained selection pressures driven by environmental change and thereby evolve adaptive phenotypic plasticity (Schlichting and Pigliucci 1998; Gonzalez et al. 2013). The presence of genetic variation in phenotypic plasticity is recorded by the presence of significant genotype-by-environment interactions (GEI:s)(Schlichting and Pigliucci 1998; Lande 2009). If a population lacks such genetic variation in reaction norm slopes, microevolutionary changes in plasticity will be prevented (Sisodia & Singh 2010; Husby *et al*. 2010).

Hansen and Houle (2004) argued that the vast majority phenotypic traits have large amounts additive genetic variation, even when such traits show evidence of long term stasis in the fossil record or in extant populations. However, and in contrast to this view, there are some documented empirical examples where genetic variation for physiological traits has been demonstrated to be low enough to act as an evolutionary limit (Blows & Hoffmann 2005). If a population is invariant in its evolutionary response to sustained environmental change, any change outside of a critical range should rapidly lead to population decline and ultimately extinction (Charmantier *et al*. 2008; Visser 2008; Chevin *et al*. 2010).

Thermal plasticity is a form of phenotypic plasticity that is particularly important for ectotherms, which have limited ability to buffer themselves against external temperature changes (Angilletta *et al*. 2002; Angiletta 2009). Many ectotherm species might already be close to their upper physiological thermal limits (Addo-Bediako *et al*. 2000; Deutsch *et al*. 2008; Kellermann *et al*. 2012). In particular, small insects and other ectotherms may lack the ability to buffer themselves against external temperatures altogether (Stevenson 1985). This might be reflected as canalization (i.e. low genetic variance in thermal reaction norms) resulting in low thermal plasticity (Angilletta *et al*. 2002; Charmantier *et al*. 2008). In general, we know relatively little about the amount of genetic variation in thermal plasticity in natural populations. Moreover, most previous studies on thermal adaptation and thermal plasticity in insects and other ectotherms focus on how temperature affects survival and hence the implications for natural selection. In contrast, the consequences of temperature challenges for sexual selection (e.g. how mating rates and mating success is affected by temperature and thermal plasticity) has seldom been a focus for empirical investigations (see Olsson et al. 2011 and Taylor et al. 2015 for exceptions). Only recently has temperature also been linked to several aspects of sexual selection and speciation in ectotherms. Examples of such recent studies exploring the link between thermal adaptation and sexual selection include how melanin-based dark colouration affects body temperatures within local populations and across latitudinal clines (Punzalan et al. 2008; Svensson and Waller 2013), how different microclimatic environments reduces immigrant male viability (Gosden et al. 2015), how mating rates might be temperature-dependent (Olsson et al. 2011; Taylor et al. 2015) and a recent finding that postzygotic isolation evolves more rapidly between various species of *Drosophila* in hot tropical areas, compared to cooler temperate areas (Yukilevich 2013).

Isogenic *Drosophila melanogaster* lines offer an excellent opportunity to investigate genetic variation in thermal reaction norms. The Drosophila Genetic Reference Panel (DGRP) is a public resource consisting of more than 200 inbred lines derived from a population in Raleigh, North Carolina (Mackay *et al*. 2012). Since the lines are isogenic, any phenotypic differences between these lines can be attributed to genetic effects, provided that they are raised and kept under identical conditions. Here we investigate and quantify the amount of genetic variation in thermal reaction norms of 30 DGRP lines, using a hemiclonal experimental approach (Abbott & Morrow 2011), and also by comparing these different DGRP-lines directly with each other. These DGRP lines should be representative sample and a snapshot of naturally segregating genetic variation in the local populations from which they were derived (Mackay *et al*. 2012) and have also been previously used them in a study on sexual selection on wing interference patterns (WIPs)(Katayama et al. 2014).

A necessary condition for the evolution of adaptive phenotypic plasticity in a novel thermal environment is that the population harbors enough standing genetic variation in the trait of interest (Chevin et al. 2010). A relatively low amount of genetic variation would indicate that the population is unlikely to respond evolutionarily to these novel thermal conditions. Here we investigate how two behavioral traits important to male fitness – male locomotion and mating success – are influenced by varying thermal conditions and whether these two fitness-related traits show any evidence for genetic variation in plasticity. We used male locomotor activity as a measure of performance as this trait is likely to be associated with fitness because of its links with reproductive success, dispersal, predator avoidance, and foraging (Gilchrist, 1996; Roberts et al., 2003; Long & Rice, 2007; Latimer et al. 2011). We complemented these analyses of behavioral performance traits with an analysis of a physiological trait – heating rate – using the technique of thermal imaging (“infrared camera”) on a subset of these DGRP-lines. Heating rate is also likely to covary with physiological performance and mating success, especially in ectotherms. Thermal imaging is a technique by which body temperatures of both endotherms and ectotherms can be quantified, under laboratory, semi-natural, and natural field conditions (Tattersall et al 2009; Tattersall and Cadena 2010; Symonds and Tattersall 2010; Svensson and Waller 2013).

## Methods

### *Drosophila melanogaster* sources

Isogenic lines used in this experiment were obtained from the Drosophila melanogaster Genetic Reference Panel (DGRP) of the Bloomington Stock Centre (Mackay *et al*. 2012), which were created after 20 generations of full sibling inbreeding of the stock inbred fly populations (Mackay et al. 2012). Wild type (LHm) flies were originally obtained from Edward H. Morrow (EHM), University of Sussex, Falmer, UK, and maintained in Lund since 2012 in the laboratory of J. Abbott. These LHm flies originated from 400 flies collected by L Harshman in central California in 1991 and they have been maintained since that time by L Harshman (1991–1995), WR Rice (1995–2004) and EHM (2004–present) (Carter et al. 2009).

### Producing hemiclonal males from DGRP-lines

A total of 16 male DGRP flies for each isogenic line were crossed with 16 wildtype virgin LHm females (in a total of 992 vials). The outbred male offspring were then used for the mating and locomotion assays. We refer to these outbred male flies as hemi-clones, following previous terminology (Abbott and Morrow 2011). This outbreeding procedure was conducted to reduce any inbreeding effects on mating behaviour that could potentially remain among the DGRP-lines (Huang et al. 2012). Moreover, by comparing male mating success in the DGRP background vs. mating success in a hemi-clonal background, we were also able to reduce the effects of non-heritable genetic variation across the different genetic backgrounds. Additive genetic variance in male mating success is expected to produce a significant correlation in male mating success between the DGRP-lines and the corresponding male genotype in the hemi-clonal background. Conversely, a weak correlation between male mating success in these different genetic backgrounds would imply low additive genetic variance for this trait and might also indicate a large non-additive effects, arising from e.g. epistatic genetic variance (Meffert et al. 2002) or dominance variation (Merilä & Sheldon 1999). All flies were cultured in a 20mm medium of cornmeal, yeast, and molasses and kept at 25°C in an incubator, on a light-dark cycle of 12hrs:12hrs.

### DGRP-lines and hemiclonal experiments

Two complementary groups of flies were used: 1) 32 pure isogenic DGRP lines, and 2) 30 hemi-clonal DGRP lines (Mackay et al. 2012). Throughout this paper, we refer to these separate experimental fly categories as DGRP-lines and hemiclonal lines, respectively. For the 32 pure DGRP-lines, the mating assay was performed in three test temperatures: 18°C (cold), 24°C (optimal) and 30°C (hot). We performed three replicates for each line and temperature treatments for these pure DGRP-lines for a total of 265 vials.

In a first pilot study using hemiclonal males, we selected 10 DGRP lines for outbreeding to produce 10 groups of hemiclonal males. These 10 DGRP lines were specifically selected for outbreeding because they showed variable patterns in their mating rates to different temperatures in the initial mating assay with the 32 pure DGRP-lines. In general, however, the individual patterns we observed in the pure DGRP lines were only weakly related to the patterns in the outbred lines (Fig. 4) (Huang et al. 2012). Hemi-clones were produced by mating 992 vials of 16 DGRP males with 16 virgin females from the outbred LH_M_ population (Chippindale et al. 2001). Male offspring from these crosses will therefore have one set of DGRP autosomes and the Y in a random LH_M_ background. For three test temperatures, 18°C (cold), 24°C (optimal) and 30°C (hot), we performed both mating and locomotion assays.

We followed up the first pilot hemiclonal study with a second study, where we used 30 isogenic lines (DGRP) to produce a new set of 30 hemiclonal groups of males. In this follow-up study, we performed mating and locomotion assays at 18°C (cold), 24°C (optimal), 30°C (hot) and we also added one additional temperature treatment at 36°C (extremely hot) (Trotta *et al*. 2006). This temperature range is similar to what has been used in other studies (Latimer et al. 2011). We performed six replicates for each line and temperature treatment in this assay, although the total number of replicates will be slightly greater for those lines in which the pilot study was also included. We pooled the results from first hemiclonal study with the second to increase statistical power, while accounting for the effect of experimental sessions as a block in our statistical analyses.

### Mating assays

Each line was anaesthetised with CO_2_ gas. Seven males per vial were collected and placed in separate 25 x 95 mm vials (with fly medium). Seven LHm virgin females were also collected per vial, and placed in separate vials. Vials of males and females were kept at 25°C overnight, to allow recovery from anaesthesia. For the experiment, vials containing hemiclonal males (one vial per line), and vials of LHm females were placed in an incubator at the test temperature and allowed to acclimatise for 30 minutes. After this time, the flies from one female vial were transferred without anaesthesia (“flipped”) into a male vial. Vials with males and females were shaken lightly to avoid early mating while the other vials were being combined. The vials, now with 14 flies in total, were placed back in the incubator at the test temperature for 1 hour as the mating assay was conducted. One vial for each of our clonal lines (hemi or pure) was placed in the incubator at a time. For each of the test temperatures, the number of copulating pairs was used a measure of male mating success. The number of mating males in each vial could thus vary from 0 to 7, and it was recorded every 10 minutes (or 15 minutes for the pure lines) over a one hour period. Here, we are measuring a mating rate, which also captures any variation in the latency to mate. All observations were recorded each day between 9.00 and 17.00hrs. We randomized the time of day for mating observations within each line.

### Locomotion assay

After 30 minute period of incubation, and before the flies from the male and female vials were combined, the male vials were tapped to cause all the males to fall to the bottom of the food vial. We then recorded the time required for the fastest male fly to walk up the side of the vial from the bottom of the food to the top of the vial plug (95 mm). This was repeated three times for each line and we took the average value per vial (Gibert *et al*. 2001).

### Heating rate assay

Using a thermal imaging camera (NEC Avio Infrared Technologies H2640) and macro lens (NEC Avio Infrared Technologies TH92-486), we recorded the heating rate of 10 DGRP lines (see supplement for information about the specific DGRP lines used). Fly individuals from each line were cooled in a climate chamber at 5°C for 3 minutes before being removed from the chamber and allowed to heat up to room temperature (approximately 23°C) in a petri dish. Thermal images were taken every 5 seconds for around 30 seconds or when all the flies had left the petri dish. Two experimental blocks of each line were performed. We recorded how body temperature changed for each line over this time period by analyzing these thermal images using the software provided by NEC Avio (see Svensson and Waller 2013 for more methodological details).

### Statistical analyses

All statistical analysis in this paper were conducted in R (R Development Core Team 2008). R-code for all the analyses are uploaded to Dryad, and details are provided in the Supplementary Material. To analyze the heating rates of 10 chosen DGRP lines, we performed an analysis of variance with temperature at each time point as the dependent variable. The following model was used: Temp = Time + Line + Block + Time*Line + Time*Block + Temp*Line*Block.

For the hemi-clones, we performed a two-way analysis of variance (ANOVA) with the number of matings at 10 minutes as the dependent variable, with line, temperature, and their interactions as dependent factors. In this analysis, line and temperature were both treated as categorical factors. In a follow-up analysis of these hemi-clonal males, we instead treated temperature as continuous variable, both as a simple linear term and as a quadratic term and their two-way interactions with line (i.e. mating rate = Temp + Line + Temp*Line + Temp^2^ + Line*Temp^2^). We chose to analyse temperature as both a continuous and categorical variable because both analyses have useful interpretative value. In the analysis of the pure DGRP-lines, we performed a similar two-way analysis of variance (ANOVA) with the number of matings at 15 minutes as the dependent variable and line, temperature and their interaction as independent dependent factors. In this analysis, line and temperature were also both treated as categorical factors.

We additionally performed a repeated measures analysis on the hemiclonal lines using the R-package ‘nlme’ (Pinheiro et al. 2016). We used the number of matings as the dependent variable, and experimental temperature category (18, 24, 30, 36 °C as different levels), line (the 30 hemi-clonal lines as different levels), and time (10, 20, 30, 40, 50, 60 minutes) were treated as fixed effects in this model. Vial and experimental block were treated as nested a random effects. This allowed us to control for the non-independence of the repeated mating counts over each vial.

The two-way interaction between line and temperature treatment in these tests, should reflect the magnitude and possible statistical significance of genotype-by-environment interaction with respect to thermal plasticity for male mating rate. Significant line-by-temperature interactions would thus indicate the presence of genetic variation in the thermal reaction norms of males belonging to different DGRP- and hemi-clones respond in terms of their mating success at different temperatures.

Finally, we performed a power analysis simulation to quantify the minimum amount of genetic variation in thermal reaction norms that we would be able to detect an effect in the 30 hemi-clonal lines, and using a given sample size (R-code for this simulation will be provided on Dryad). To analysis variation in heating rate, we performed a two-way analysis of variance (ANOVA) with the average temperature at each time point as the dependent variable and line, time, and their interactions as dependent factors, while also controlling for a block effect. An analysis modelling mating rate as a proportion is presented in the Supplementary Material (Table S1).

## Results

We found significant variation in mating rates and locomotory performance among lines, using both the hemi-clonal and the pure DGRP lines (Tables 1-4, S1, S2, S3; Fig. 1). As expected, all lines of both hemi-clonal and pure DGRP experimental categories responded plastically to the different temperature treatments (Fig. 2). Male mating rates and locomotory performance were significantly affected by temperature (Fig 2).

**Table 1.** Analysis of variance (ANOVA) for the mating rate in hemi-clonal males. Line (N=30) and temperature (N=4; 18,24,30,36 °C) were both treated as categorical factors. We used the first mating observation at 10 minutes to avoid double counting matings at the next observation time 20 minutes. Here, a significant Line x Temp interaction would be indicative of a GEI and reveal significant genetic variation (greater than zero) in plasticity in our hemiclonal males. Block is a categorical factor (N=2), which controls for differences in the two experimental runs. See table S1 in the supplementary material and a model wherein mating rate is treated as a proportion.

| Term | DF | SSE | MSE | F-value | P-value | Significance |
| --- | --- | --- | --- | --- | --- | --- |
| Line | 29 | 4.54 | 0.16 | 2.37 | <0.0001 | **** |
| Temp | 3 | 2.48 | 0.83 | 12.51 | <0.0001 | **** |
| Block | 1 | 0.3 | 0.3 | 4.52 | 0.034 | * |
| Line x Temp | 87 | 4.88 | 0.06 | 0.85 | 0.83 | ns |
| Residuals | 870 | 57.38 | 0.07 |  |  |  |

**Table 2.** Analysis of variance (ANOVA) for the mating rate of the DGRP males. Line (N=32) and temperature (N=3; 18,24,30 °C) were both treated as categorical factors. We used the first observation time at 15 minutes to avoid double counting matings at the next observation time 30 minutes. A significant Line x Temp interaction would be indicative of a GEI.

| Term | DF | SSE | MSE | F-value | P-value | Significance |
| --- | --- | --- | --- | --- | --- | --- |
| Line | 31 | 5.22 | 0.17 | 3.55 | <0.0001 | **** |
| Temp | 2 | 1.31 | 0.66 | 13.82 | <0.0001 | **** |
| Line x Temp | 58 | 2.35 | 0.04 | 0.86 | 0.752 | ns |
| Residuals | 172 | 8.16 | 0.05 |  |  |  |

**Table 3.**
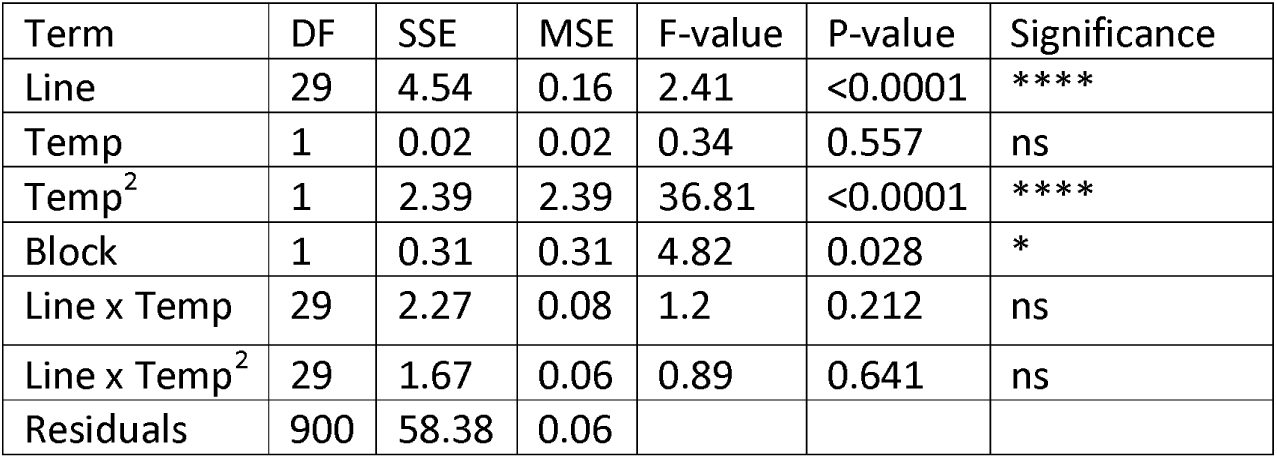
Analysis of covariance (ANOVA) for the mating rate hemi-clonal males. Here Line (N=30) is treated as a categorical factor and temperature is treated as a continuous variable. The quadratic effect of temperature (Temp^2^) allows the model to detect any curvature in the reaction of mating rate to temperature. We used the first observation time at 10 minutes to avoid double counting matings at the next observation time 20 minutes. Here, a significant Line x Temp interaction would be indicative of a GEI, and indicate significant genetic variation (greater than zero) in plasticity in our hemi-clonal males. Block is a categorical factor (N=2), which controls for differences in the two experimental runs.

**Table 4.** Analysis of locomotion performance in the hemi-clonal lines (N=30). Locomotion is the speed (mm/seconds) of the fastest hemiclonal male (7 per vial) to walk up the side of the vial (repeated 3 times for each line) at the different experimental temperatures. We performed a two-way analysis of variance (ANOVA) with locomotion time the as the dependent variable and line, temperature and their interaction as independent dependent factors.

| Term | DF | SSE | MSE | F-value | P-value | Significance |
| --- | --- | --- | --- | --- | --- | --- |
| Line | 29 | 4197.69 | 144.75 | 1.76 | 0.009 | ** |
| Temp | 3 | 7449.08 | 2483.03 | 30.22 | <0.0001 | **** |
| Line x Temp | 87 | 5691.16 | 65.42 | 0.8 | 0.907 | ns |
| Residuals | 592 | 48635.14 | 82.15 |  |  |  |

**Fig 1.**
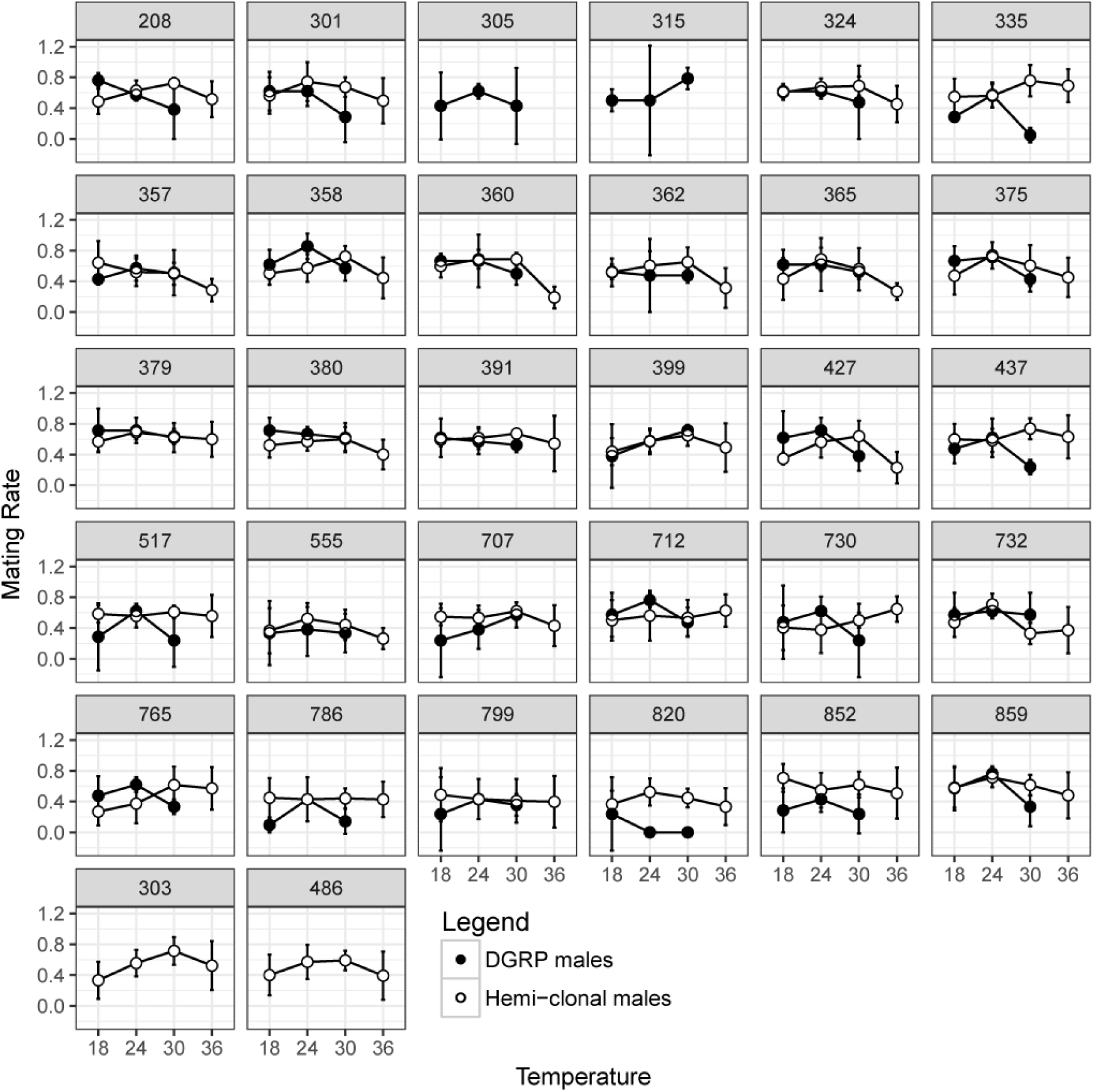
Thermal reaction norms of male mating success estimated for of each pure DGRP (black) and hemi-clonal (empty) DGRP-line in relation to experimental temperature treatments. Numbers above each sub-panel refer to the DGRP-lines described in Mackay et al. (2012). Although temperature affects mating rates in all lines and often peaks at intermediate temperatures, there is no evidence for any genotype-by-environment interaction with respect to temperature, i.e. the mating success of the different lines are similarly affected across environmental treatments. Mating rate is the number of copulations that were recorded in a vial after 10 minutes (for the hemiclones) and after 15 minutes (for the pure clones). The mating assay was only performed for some lines with either only the hemi-clones (303, 486) or only the pure lines (305, 315). Confidence intervals (95%) are shown as vertical lines.

**Fig 2.**
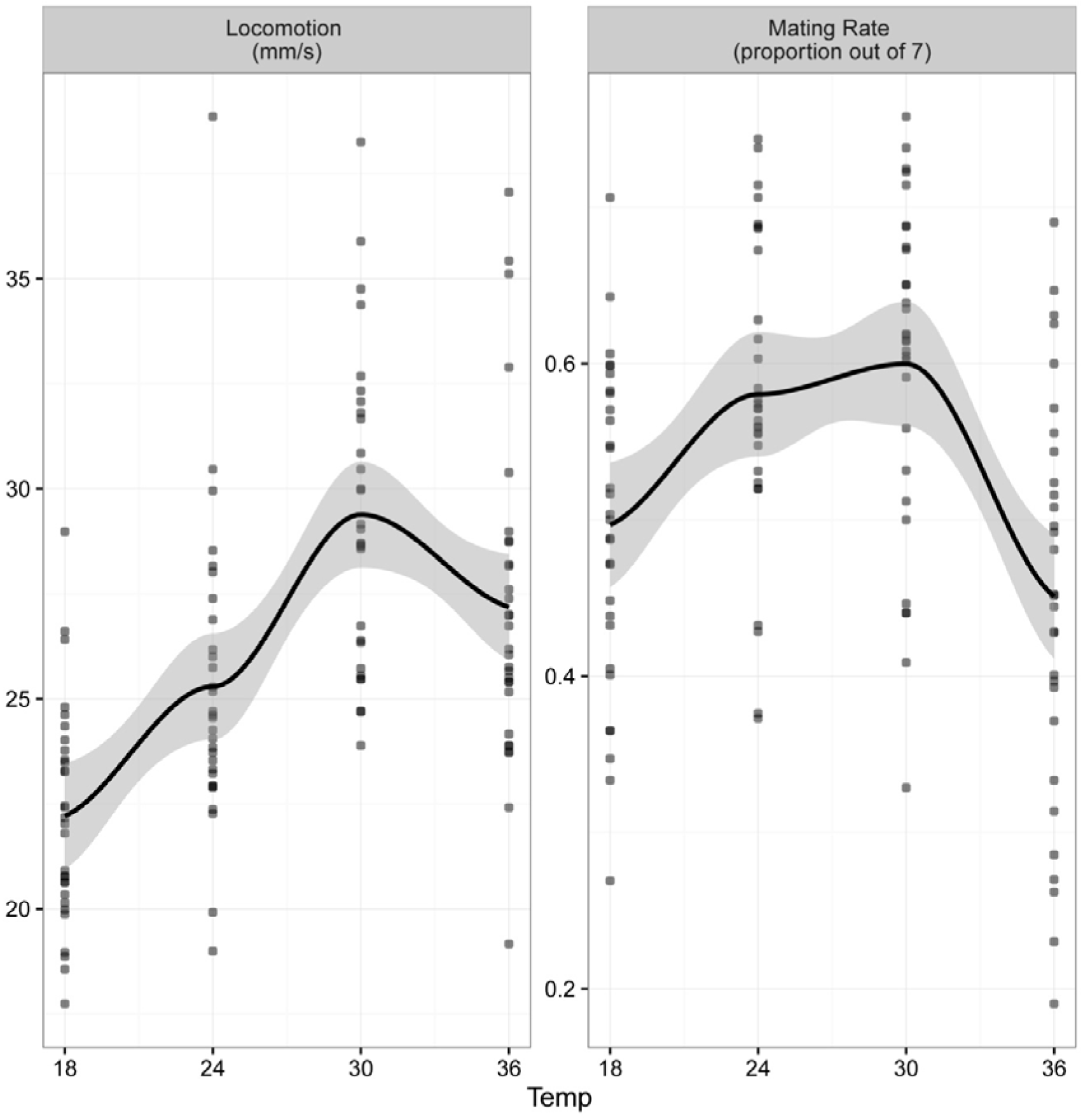
Temperature-related male performance for the hemi-clonal males showed as male locomotory performance (left panel) and male mating rate (right panel). Males from different lines vary in mean mating rates and locomotory performance (intercepts), but variation between lines in thermal reaction norms (slopes) is low (Fig. 1). Hence, there is no evidence for significant genetic variation in the thermal reaction norms (Fig. 1). Each data point represents the mean performance of each line at each temperature treatment.

For all of our statistical analyses, and for neither the hemi-clonal nor the pure DGRP assays, we did not find any statistically significant interaction between line and temperature, that would be indicative of GEI:s (Fig. 1; Tables 1-3, S1, S2, S3). Moreover, we were not able to find any evidence for a statistically significant interaction between line and the squared temperature component (Table 3), i.e. neither the slopes nor the curvatures of the thermal response curves differed significantly between lines (Fig. 1).

Visual inspection of the thermal performance curves for male mating success in Fig. 1 revealed that 24 out of the 30 hemi-clonal lines peaked at intermediate temperatures (24 °C or 30 °C), whereas only 6 lines had maximal mating success in the cold (18 °C) or extremely hot treatments (36 °C). For the DGRP-lines, 17 peaked in male mating success at intermediate temperature (24 °C), 9 at cold temperature (18 °C) and the remaining four at 30 °C (note that the DGRP-lines were not evaluated under the extreme treatment (36 °C; see Fig. 1). However, note that the relationship between male mating success and temperature was often quite flat in the range between 18 and 24 °C, after which it dropped (Fig. 1). Taken together, these results suggest a genetically quite invariant intermediate temperature optimum and more or less flat fitness peak around 24 °C (Fig. 2). Thus, we found no evidence for any statistically significant difference between genotypes in the location of this fitness optimum and neither any evidence for different slopes of the thermal response curves or their curvatures (Table 3).

Using our statistical power simulation we were able to put a minimum bound on the genetic variation in thermal plasticity (variation in the effect of our Line x Temp interaction) (Fig. 3). The main conclusion from these simulations is that our statistical power is high enough under realistic parameter values, meaning that we should have detected a large GEI:s if they had existed (Fig. 3). The magnitude of the GEI we recorded in these experiments can explain at most 4% of variation in mating rate. For instance, our statistical power approaches one with an effect size of 0.03 (treating temperature as a continuous variable) and 0.15 (treating temperature as a categorical variable)(Fig. 3). Finally, at 18° and 30°C there was no detectable correlations between the mating rate of the hemi-clones and our DGRP lines. At 24° C, there was a slightly stronger and significant correlation between the mating rates of the DGRP and hemi-clonal males (Fig. 4).

**Fig 3.**
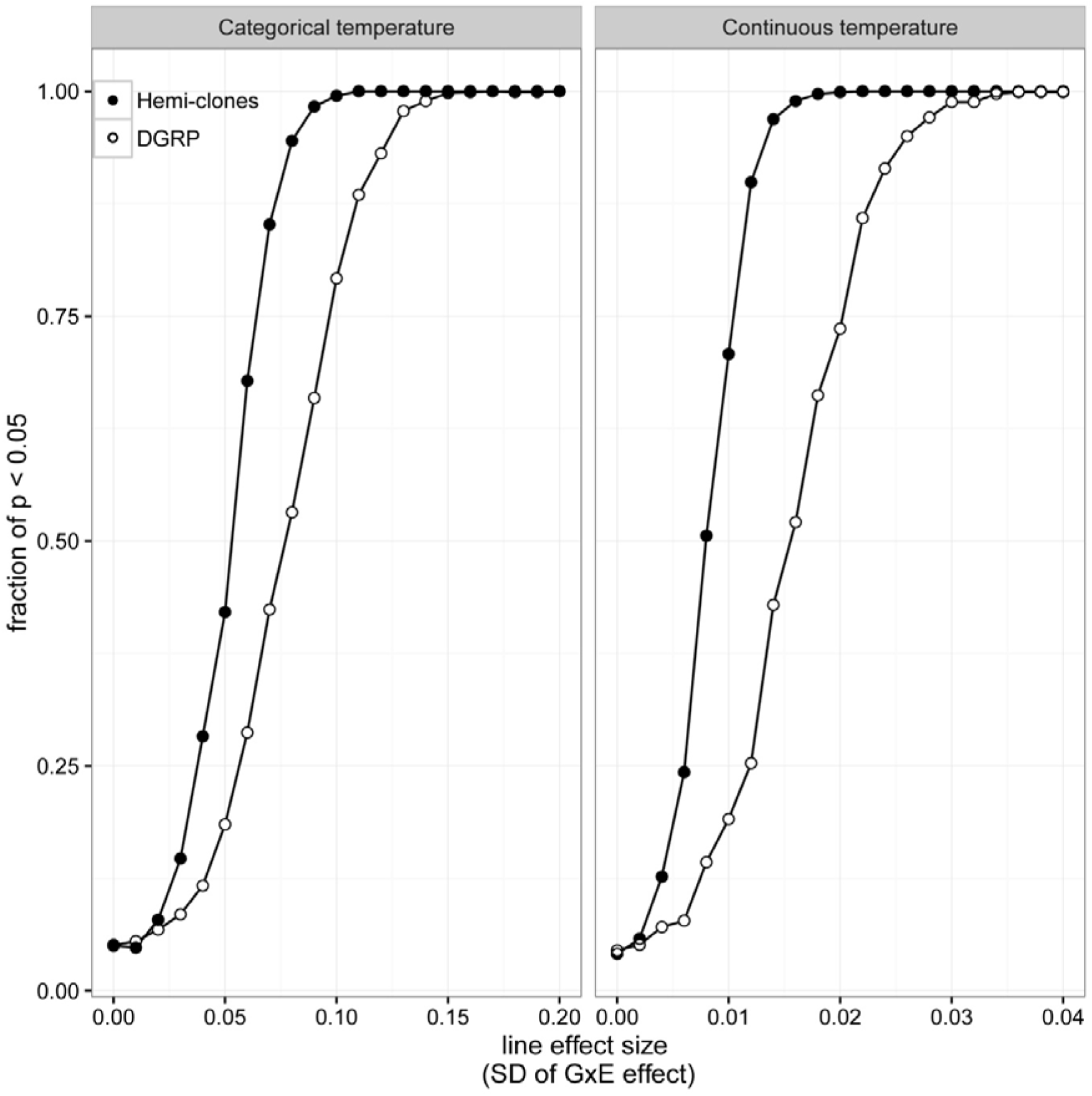
Power plots to detect significant GEI:s (reflected as Line*Temp interactions) for male mating success. Temperature is either treated as continuous variable (left plot) or as a discrete experimental factor (right plot). Comparing the slopes of the reaction norm (left panel) from optimal temperature to extreme (hemiclonal lines: 24° to 36° C; pure DGRP-lines: 24° to 30 ° C), we found that the standard deviation in slope is likely to be < 0.01 for the hemi-clonal lines and < 0.025 for the pure DGRP-lines. This means that lines varied in less ±0.008 matings (per 10 min) per 1° C. Treating temperature as discrete (right panel), the standard deviation of this Line*Temp effect is most likely < 0.07 for the hemi-clonal lines and < 0.12 for the pure clonal lines. This means that the variation in mating rate is maximally < 0.01 matings for a given temperature. These are minimum effect sizes, and the true variation in thermal plasticity is likely to be lower. R-code for these power simulations are provided on Dryad. Each data point represents 1000 simulations (hemi-clonal: n = 7 and pure-clonal: n = 3 replicates per line n = 30).

**Fig 4.**
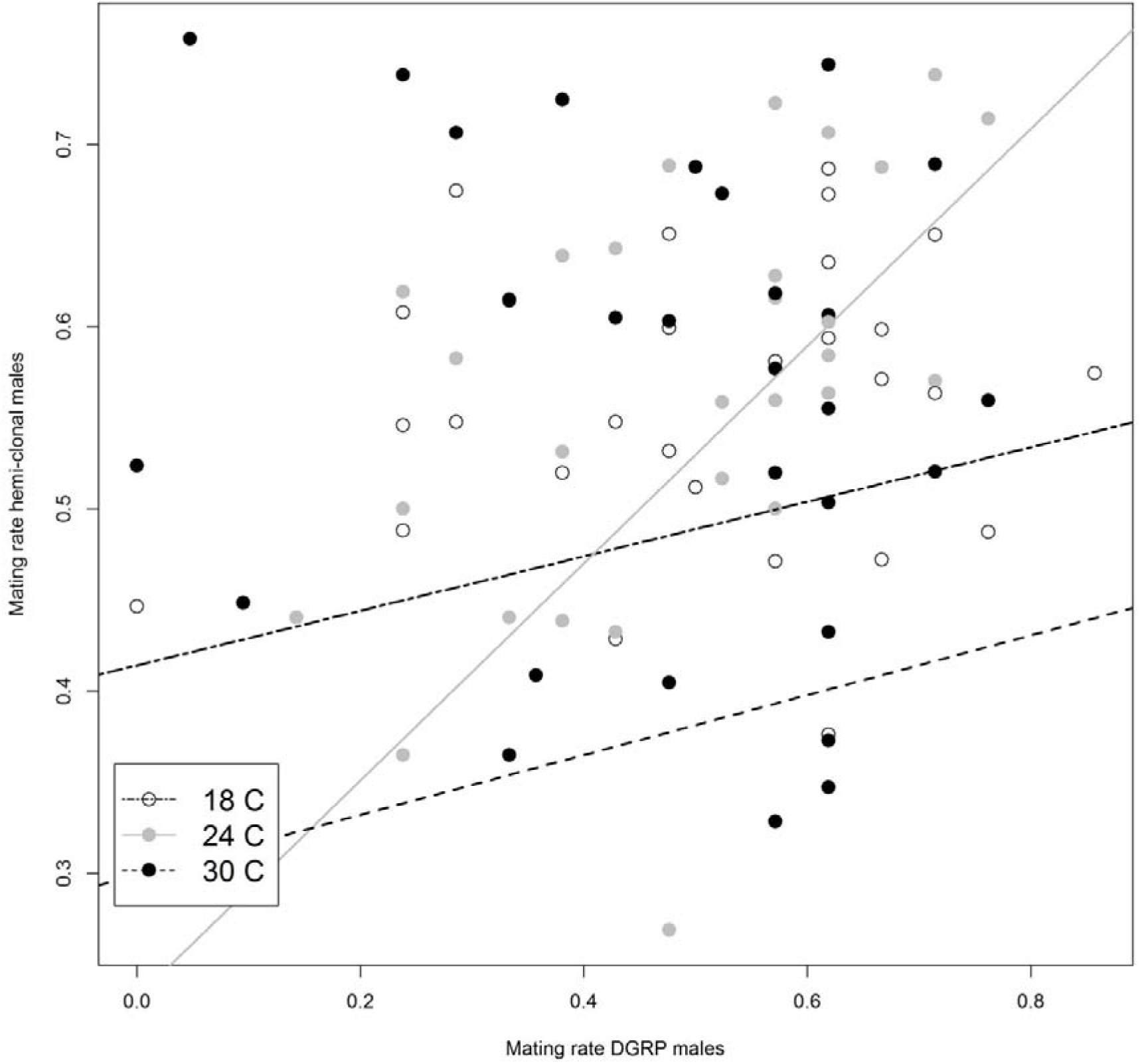
Correlations between mating rates (at 10 min and 15 between our hemi-clonal and DGRP lines, measured at three different temperatures (measured at 18 °C (“cold” = white), 24 °C (“optimal” = gray) and 30 °C (“hot” = black). We studied a total of 28 Lines per temperature treatment, i.e. the total number of datapoints in this graph is 84). The “random” Lhm genetic background in our hemi-clonal lines likely has a large effect on mating rate, since the mating rates the DGRP-lines, put either in to hemi-clones or measured as pure clones show only weak non-significant relationships (All: r=0.12, *P*=0.24, N=84, 18C: r=0.08, *P* = 0.68, N=29; 24C: r= 0.37, *P*=0.05, N=29; 30C: r= 0.10, *P*=0.61, N=29). The lack of strong significant correlations between the mating rates of the different lines in different genetic backgrounds indicates low additive genetic variance in male mating success and implies strong epistatic effects on mating rate (Huang et al. 2012), as our hemi-clonal and pure-clonal lines will only share the additive effects of their genotypes.

In the analysis of heating rates using thermal imaging, we found significant variation among the 10 DGRP-lines we investigated, and a significant Line x Time interaction (Fig. 5; Table 5). This shows that for these 10 DGRP-lines, the slopes of the thermal reaction norms differed, although the effect was significant only in the first experimental block (Fig. 5; Table 5).

**Fig 5.**
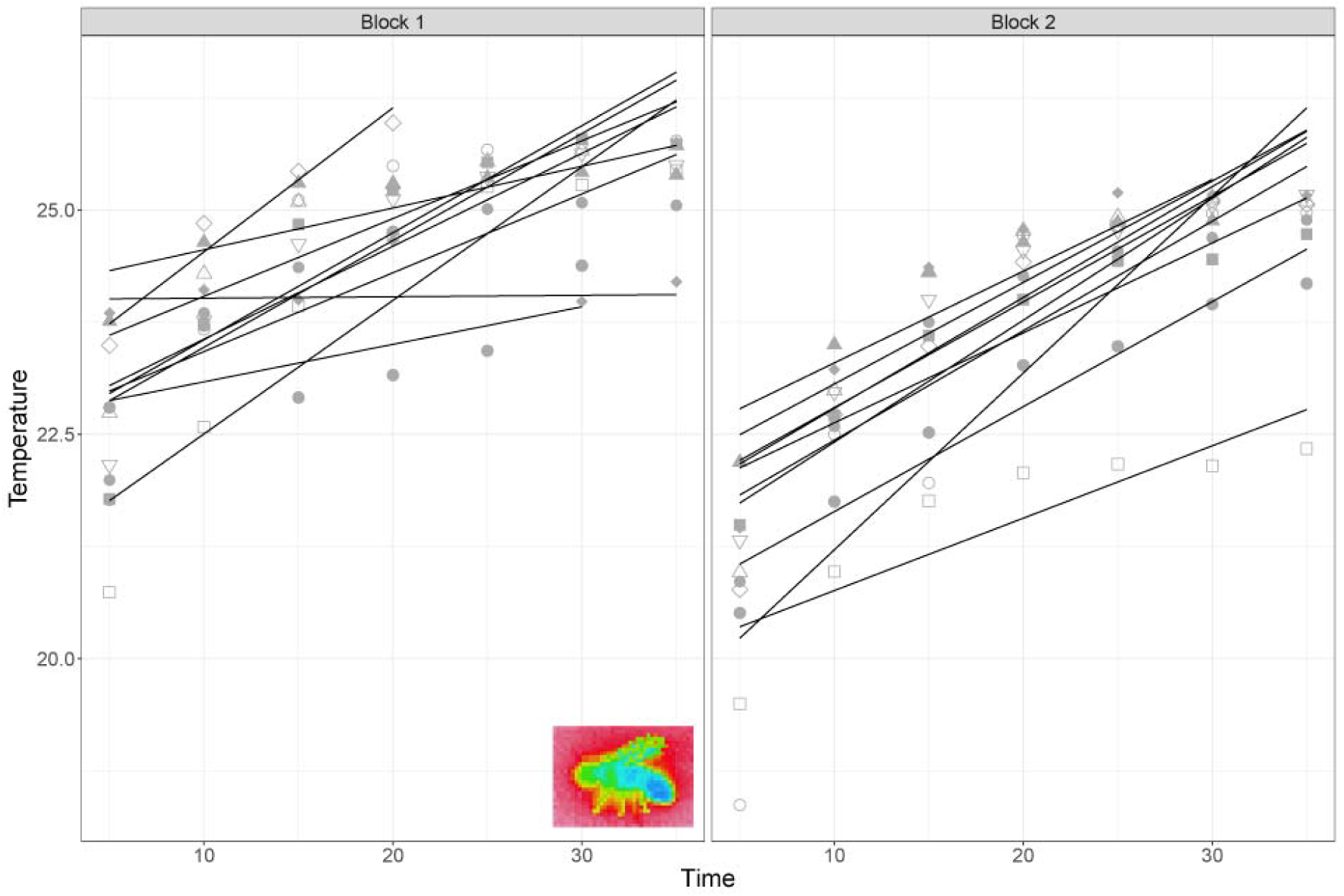
**A)** Heating rates of 10 DGRP lines after exposer to a cool shock 5°C for 3 minutes and then allowed to recover at room temperature for 30 seconds. Two experimental blocks were performed of these same 10 DGRP lines. Thermal images were taken every 5 seconds. Heating rate was significantly variable between lines in block 1 but not in block 2 (Table 5).

**Table 5.** Analysis of variance (ANOVA) for the heating rate of DGRP males. 2 replicates of Lines (N=10). Time is measured in seconds. A thermal image was taken at each time point. Here, a significant Line x Time interaction would be indicative of significant variation in heating rate between lines. However, a significant Time x Line x Block effect complicates this interpretation. Block is a categorical factor (N=2), which controls for differences in the two experimental runs.

| Term | DF | SSE | MSE | F-value | P-value | Significance |
| --- | --- | --- | --- | --- | --- | --- |
| Time | 1 | 135.14 | 135.14 | 290.3 | <0.0001 | *** |
| Line | 9 | 42.62 | 4.74 | 10.17 | <0.0001 | *** |
| Block | 1 | 35.44 | 35.44 | 76.13 | <0.0001 | *** |
| Time x Line | 9 | 9.46 | 1.05 | 2.26 | 0.024 | * |
| Time x Block | 1 | 3.94 | 3.94 | 8.47 | 0.004 | ** |
| Line x Block | 9 | 16.48 | 1.83 | 3.93 | 0.0002 | *** |
| Time x Line x Block | 9 | 8.47 | 0.94 | 2.02 | 0.04 | * |
| Residuals | 95 | 44.22 | 0.47 |  |  |  |

## Discussion

Evolutionary change requires genetic variation in the traits under selection (Blows & Hoffmann 2005). This basic requirement for evolutionary change also applies to a trait like thermal plasticity, and the evolution of thermal plasticity will require genetic variation in the slopes of thermal reaction norms. In this study, we have found significant genetic variation in male mating rates, using two different approaches: a pure-clonal approach (using a subset of un-manipulated DGRP-lines bred by Mackay et al. 2012) and a hemi-clonal approach (Abbott and Morrow 2011), where these DGRP-lines were introduced into an outbred LH_M_ background (Fig. 1). Using these two complementary experimental approaches, we found no statistically significant variation in the reaction norm of male mating rate to temperature in the different lines (Figs. 1, 3). This shows that although all these lines altered their mating rates in relation to temperature (i.e. phenotypic plasticity; Fig. 1, Table 1-4) the changes were parallel and all lines responded in a similar manner. This implies that genetic variation in thermal plasticity for male mating success is low (Fig. 1), explaining at most 4 % of the variation (Fig. 3). Similar conclusions apply to male locomotory performance (Fig. 2), where we also did not find any evidence for significant GEI (Table 4).

The lack of strong GEI:s for these two behavioural traits contrasts our findings of a significant GEI for male heating rate, a physiological performance trait, measured using thermal imaging (Fig. 5), where we did find evidence for GEI (Table 5). Heating rate is an important fitness trait for ectotherms because it might covary with ability to adjust to natural ambient temperatures. It also measures the capacity of an individual to buffer itself against the ambient environmental temperatures. The use of thermal imaging is a very powerful tool to quantify genetic and phenotypic variation of a physiological trait like heating rate, as done in previous studies of non-model organisms (Tattersall et al 2009; Tattersall and Cadena 2010; Symonds and Tattersall 2010; Svensson and Waller 2013). The fact that the significant GEI for thermal reaction norm slopes in these heating rates have no counterpart in the behavioural assays of mating rates and locomotory assays (Figs. 1-2; Tables 1-4) might be biologically important. Behavioural traits and life-history traits are further downstream than the physiological traits are from the genes that govern phenotypic traits (Price and Schluter 1991), hence genetic variation on these grounds expected to be lower for such higher-level traits, such as mating rate and locomotory performance.

Overall, and across all lines, male mating success was maximal at 24 °C (DGRP-lines) or at either 24 °C or 30 °C (Fig. 2), with a few exceptions (Fig. 1). This suggests that the thermal optimum for male mating success falls well within the normal temperature range *Drosophila melanogaster* will experience in North Carolina in the wild (Annual high temperature: 21.5°, Annual low temperature: 9.3°, Average temperature: 14.9° C (Daly 2000)), where these DGRP-lines originated (Mackay et al. 2012). Thus, although our results suggests limited evolutionary potential for thermal plasticity with respect to male mating success, they are consistent with males being locally adapted with respect to their local temperature regime, consistent with a previous study on *Drosophila melanogaster* (Dolgin et al. 2006, Latimer et al. 2011).

These DGRP lines were derived from field-caught flies and variation among these lines should reflect naturally segregating genetic variation in the source population (Mackay *et al*. 2012). Using a statistical power simulation (Fig. 3), we were able to put a minimum bound on the amount of variation in reaction norms in our clonal lines. The line effect had a standard deviation of < 0.5, which implies that variation in mating rate was at a maximum less than 0.01 matings for any given temperature. Comparing the slopes of the reaction norms from optimal temperature condition (24° C) to extreme temperature condition (36° C), we conclude that the standard deviation in slopes is likely < 0.01. This effect covers only about 4% of the total variation in male mating success (Line = 11%, Temp = 58%, Block = 21%, Line:Temp = 4%, Residual = 5%). This means that lines varied less than around ±0.008 matings (in 10 min) per 1° C (Fig. 3). Whether this low amount of variation in thermal reaction norms would allow for evolution of adaptive phenotypic plasticity is an open question and depends on several other ecological and evolutionary factors, including population size, the strength of selection, the rate of environmental change, generation time, and intrinsic rate of increase (Hoffmann 2010; Chevin *et al*. 2010). However, we note that the effect sizes in these power calculations are minimum effect sizes, and the true amount of genetic variation in thermal plasticity might be considerably lower.

One concern is that mating rate is the product of the behaviour of multiple individuals interacting, so it is not only the male’s behaviour that matters, but also the female’s preference. For instance, LHm females prefer males from some of the DGRP-lines more than males from other DGRP-lines. A second concern is that it is the additive genetic variation that determines the evolutionary potential of a population, yet the hemi-clones still share the same half-genome, so any epistatic effects arising from interactions between chromosomes within the DGRP half will also be included. Our experimental design is for these reasons conservative with respect to our ability to detect significant GEI:s, since the line-effects will partly also include non-additive effects. Additionally, there might exist variation in latency to mate after a disturbance between lines, and this might be of some concern. However, any differences in willingness to mate after disturbance will be captured by the line effect in our statistical analyses. This means that lines might differ not only mating rate per se but also in their willingness to mate at each temperature. This variation in willingness to mate after a disturbance would not interfere with detecting GEI:s in mating rate, unless there was an interaction between the latency to mate and temperature treatment.

For the hemi-clonal lines, epistatic effects would not be included by the line factor, since our starting iso-genetic lines (30) were homozygous at all loci. Thus, epistatic interactions with the outbred (LHm) genetic background would not be included in the line effect, and would become part of the error variance. Such epistatic genetic variance would represent hidden genetic variation that we were not able to detect with our hemi-clonal lines (Huang *et al*. 2012; Mackay 2014). Epistatic variance of traits related to mating success, such as courtship, have been demonstrated for other species of flies (Meffert et al. 2002). In these previous studies it has been shown that such epistatic variance can be converted to additive genetic variance following population bottlenecks (Meffert et al. 2002). From a theoretical viewpoint, traits that are closely related to fitness such as male mating success (a major fitness component) are expected to show low additive genetic variance, due to the depleting effects of strong directional sexual selection (Rowe & Houle 1996). Directional sexual selection should therefore expect reduce the additive genetic variance fraction for male mating success, resulting in a relatively higher fraction of the remaining genetic variation being non-additive, reflecting either epistatic (Meffert et al. 2002) or dominance variance (Merilä & Sheldon 2002). Such non-additive genetic variance for male mating success could potentially explain the low concordance between male mating success in the DGRP-lines and the hemi-clonal lines (Fig. 4). This interpretation of high epistatic variance for male mating success in these DGRP-lines would be consistent with previous studies of these DGRP-lines, where high epistatic variance was found for a number of other fitness-related traits, including cold tolerance (measured as chill coma recovery) (Huang et al. 2012).

Our results have some implications for the prospects of evolutionary rescue through the evolution of adaptive phenotypic plasticity (Chevin et al. 2013) and by sexual selection (Candolin & Heuschele 2008). Previous laboratory experiments on several species of *Drosophila* (Holland 2002; Rundle et al. 2006) and seed beetles *Callosubruchus maculatus* (Martinossi-Allibert et al. 2016) have found at most weak or at best mixed support for sexual selection improving the rate of evolution of local adaptation to novel stressful environments, such as thermally challenging environments. The results in the present study add to this small but growing body of literature, and indicate that thermal plasticity for male mating success is unlikely to evolve, as the genetic variation in thermal reaction norms is limited (Fig. 3). Some theoretical models suggest that sexual selection could improve environmental adaptation at low demographic costs, due to purging of deleterious alleles in males (Agrawal 2001; Siller 2001; Whitlock & Agrawal 2009). However, the results in this and other experimental studies (cited above) give only weak support to these models. Thus, our results provide only limited support to the hypothesis that sexual selection could act as an evolutionary rescuer of populations experiencing rapid environmental change (Candolin & Heuschele 2008).

Our results also agree with other research in this area showing that evolutionary responses to novel and challenging thermal conditions may be constrained (Bennett & Lenski 1993; Kellermann *et al*. 2009; Mitchell & Hoffmann 2010; Hoffmann 2010; Kelly *et al*. 2012; Kelly *et al*. 2013; Kristensen *et al*. 2015). In particular, ectotherms, particularly those living in tropical areas that already experience temperatures close to their upper thermal tolerance limits might have a reduced capacity to adapt to higher temperatures via evolutionary means (Araujo *et al*. 2013; Kristensen *et al*. 2015) (Fig 2). Insights from studies of niche conservatism also suggests that it might be difficult for many species to evolve new physiological limits (Parsons 1982; Kimura & Beppu 1993; Wiens & Graham 2005; Kellermann *et al*. 2009; Hoffmann 2010). For example, the invasive species *Drosophilia subobscura* has a range in the Americas that is restricted to climates that are similar to those found in its native range in Europe (Prevosti *et al*. 1988; Gilchrist *et al*. 2008; Huey & Pascual 2009; Hoffmann 2010). However, some natural *Drosophila* populations have adapted to their local climates (Trotta *et al*. 2006). For instance, warm-adapted populations in India have higher levels of desiccation resistance and melanism compared with cold-adapted populations (Parkash *et al*. 2008). Similarly, populations in Africa are more viable at warm temperatures than temperate populations (Bouletreau-Merle *et al*. 2003; David *et al*. 2004; David *et al*. 2005; Hoffmann 2010). Other laboratory studies have shown that populations might have the capacity to adapt to temperature via evolutionary adaptation (Hoffmann 2010; Latimer et al. 2011; Mukuka *et al*. 2010; Hoffmann *et al*. 2013; van Heerwaarden and Sgro 2013; Blackburn *et al*. 2014; van Heerwaarden at al. 2016). For example, in some laboratory experiments, *Drosophilia* have been successfully selected for increased survival after cold and heat shocks (Bubliy & Loeschcke 2005). Finally, it might be that the expression of additive genetic variation for heat tolerance is contingent on developmental temperature (van Heerwaarden at al. 2016). For instance, two rainforest *Drosophila* species exhibited significant levels of additive genetic variation when raised in warmer environments, but not when raised at 25°C (van Heerwaarden at al. 2016).

## Conclusions

The capacity of natural populations of *Drosophila melanogaster* to adapt evolutionarily (i.e. via changing the genetic composition of the population) to novel thermal conditions through the evolution of adaptive phenotypic plasticity remains an open question. In this study, we were not able to detect any significant genetic variation in thermal reaction norms to different thermal environments in either the DGRP-lines or the hemi-clonal lines. However, we were able to put a lower bound on the amount of genetic variance in thermal plasticity. Isogenic lines such as these DGRP-lines in combination with hemi-clonal approaches offer opportunities to quantify genetic variation in phenotypes where the underlying genotypes are known and line-differences can attributed to genetic differences without the need for complex breeding designs (Abbott & Morrow 2011). Such isogenic lines could also be raised in different temperatures in the future, which might reveal that significant genetic variation exists, but this variation might depend on temperatures experienced during the larval development stage (van Heerwaarden at al. 2016). It is possible that GEI:s might be more pronounced among juveniles, and that strong selection during these earlier life stages might have reduced additive genetic variance in thermal reaction norms that could be detected during the adult stage (cf. Martinossi-Allibert et al. 2016). In the present study, we were not able to detect any significant genetic variation in thermal plasticity in adult male mating success or locomotion.

## Acknowledgements

We are grateful to anonymous referees for constructive criticisms earlier drafts of this manuscript. Funding for this study has been provided by the Swedish Research Council (VR; grant no. 621-2012-3768) and Gyllenstiernska Krapperupstiftelsen (grant no. 2014-0032) to E.I. S.

